# Computational Studies of Human Sodium/glucose cotransporters: (SGLT1), (SGLT2) and investigation of Human Sirtuins together with Polydatin

**DOI:** 10.1101/2023.06.01.543288

**Authors:** Ivan Vito Ferrari, Alex De Gregorio, Maria Pia Fuggetta, Giampietro Ravagnan

## Abstract

For the first time, we investigated through molecular docking analysis with Autodock Vina and Autodock 4 the Polydatin, a derivative of resveratrol, with Sodium/glucose cotransporter 2 (SGLT1) and Sodium/glucose cotransporter 2(SGLT2) and with Sirtuins proteins, reporting excellent results both in terms of binding energies scores and inhibition constant Ki. In particular, from our analyses, Polydatin appears to have an excellent energetic affinity with human SGLT2 on the one hand, and with the human Sirtuin6, even though, comparing the binding energy values with all the investigated proteins, no significant differences were found in termins of binding energies scores. An important aspect that we want to underline is that our computational analyzes, although very accurate, require investigations in Vitro, in Vivo, and clinical studies to confirm that Polydatin has a key role with SGLT2, SGLT1, and with the Sirtuin family.

## 1. Introduction

The study that we present is focused on the natural molecule called Polydatin (also named Piceid, (3,4′,5-triidrossistilbene-3-β-d-glucoside, PD). It is a natural precursor and glycoside form of resveratrol, a stilbene commonly found in foods.^1^ It is beta-D-glucoside extract of *Polygonum cuspidatum* (or *Fallopia japonica*). It is a species of herbaceous perennial plant in the knotweed and buckwheat family Polygonaceae, a family of flowering plants informally known as the knotweed family, comprising about 1200 species that are present throughout the world. *Polygonum cuspidatum*, a root of a Chinese herb is native to East Asia in Japan, China and Korea. ^1^

According to several studies in literature, this polyphenol has shown greater stability and biological activity, in particular on the cardiovascular system compared to resveratrol. ^2-8^ Indeed many works demonstrated its excellent pharmaceutical properties and its pharmacological effects. Qiao-Hui Du et al., 2013 reported its effects on heart muscle cells to protect myocardial cells (MC) from injury caused by oxygen and glucose deprivation (OGD), ^3^ as well as was also demonstrated by (Luo et al., 1990).^4^

Kim SanTang and Jey Sern Tan 2019 focused on the role of PD in cerebral ischemia, showing neuroprotective effects, according to Vitro and In Vivo studies^6,7^. Although clinical trials are needed to evaluate its therapeutic potential in humans.

Unfortunately, there are not many toxicological studies reported on this compound.^5^ Lanzilli G, et al., 2012 investigated the anti-inflammatory effect of resveratrol and polydatin by in vitro IL-17 modulation.

Their results obtained from activated mononuclear cells in human peripheral blood, stimulated with anti-CD3/anti-CD28 monoclonal antibodies and treated with these polyphenolic compounds, have shown a significant reduction of IL-17.^7^ Indeed, several papers reported that PD has anti-inflammatory and anti-oxidative stress properties.^6,7^

Wu et a., 2020 focused on PD in the treatment of atherosclerosis, primarily playing a role in three aspects: anti-inflammatory, regulating lipid metabolism, and anti-oxidative stress.

In this work, for the first time, we examined by Docking investigation different Human proteins involved with Polydatin to predict accurately its biological role, type residues interactions bond, binding energy, ligand efficiency, and inhibition constant ki.

In short, the Molecular Docking method is an In Silico method well known, focused on the study of the Binding Energies scores (kcal mol ^-1^), the types of bonds involved, and the aminoacids involved in the active site of the proteins or target receptors.

In this way, it is possible to theoretically understand which receptors or proteins are more predisposed to bind with the molecule or drug investigated. The molecular docking method has the potential to significantly improve accuracy and predictive power in drug discovery and other applications.^9-11^ In our case, we focused on the Structure of human SGLT2 (Sodium/glucose cotransporter 2) and human SGLT1 Sodium/glucose cotransporter 1) as well as investigating all human sirtuins proteins.

We have chosen to carry out these bioinformatic studies with these proteins with polydatin for two important reasons: the first is not reported to date many works on Docking on these protein targets and the second because both the Sodium/glucose cotransporters 1-2 and the Sirtuins are involved in various biological processes. Indeed, Human sodium-glucose cotransporter 2 (hSGLT2), a member of the sodium-glucose cotransporter family, is located in the early proximal tubule that mediates the reabsorption of the majority of filtered glucose in the kidney1. Pharmacological inhibition of hSGLT2 by oral small-molecule inhibitors, such as empagliflozin, leads to enhanced glucose excretion and is widely used for the treatment of type 2 diabetes. SGLT2 inhibitors or called gliflozins the treatment of type II diabetes mellitus (T2DM) have been shown benefit patients with atherosclerotic major adverse cardiovascular events (MACE). One such study defined MACE as the compound of myocardial infarction, stroke, or cardiovascular death.^12-14^

As regards the mammalian sirtuin protein family (comprising SIRT1–SIRT7) received well-known attention for its regulatory role, mainly in metabolism and aging. Sirtuins mediate phenomena such as aging, regulation of transcription, apoptosis, and resistance to stress and also affect energy efficiency and alertness during situations with low caloric intake (calorie restriction).^15-17^

Sirtuins act in different cellular compartments: they deacetylate histones and various transcriptional regulators in the nucleus, but also specific proteins in other cellular compartments, such as in the cytoplasm and in mitochondria. As a consequence, sirtuins regulate fat and glucose metabolism in response to physiological changes in energy levels, by acting as metabolic sensors.^15-17^ Several studies on sirtuins have been focused on diseases affecting the heart.^17-20^ Recently in 2023, Yang, X et al., demostrated that SIRT2 inhibition protects against cardiac hypertrophy and heart failure. The aim of this communication is to investigate polydatin by Molecular Docking approaches by studying its energetic affinities with proteins on the one hand and with Sodium/glucose cotransporter 1 and 2 on the other to understand if this natural substance may have a significant biological role.

## 2. Matherial and Methods

### Protein selection and preparation

All proteins were dowloaded from Protein Data bank (https://www.rcsb.org) and they are accurately prepared, and finally they are saved in PDB format.

The 3D proteins investigated are: 3D Structure of Sodium/glucose cotransporter 2, (PDB Code: 7VSI), and 3D Structure of Sodium/glucose cotransporter (PDB Code: 7WMV), Structure of NAD-dependent protein deacetylase SIRT1,(PDB Code: 4KXQ), Crystal structure of human SIRT2, PDB Code: 5D7P),Structure of human SIRT3 (PDB Code: 4BN4), Structure of human SIRT4,(PDB Code:5OJN),Structure of human SIRT5,(PDB Code:2NYR),Structure of human SIRT6 (PDB Code:3K35), Structure of human SIRT7,(PDB Code:5IQZ).

All proteins were prepared prior to docking as follows: UCSF Chimera software (https://www.cgl.ucsf.edu/chimera) was used to visualize, remove crystal ligands, hetatoms and keep only Chain A, then to convert sdf files to pdb format. Hydrogen bond structures were optimized and atoms in missing loops or side chains were added. Water molecules were removed and files were saved in PDB file format.

### Ligand preparation

Structure of Polydatin was downloaded from PubChem (https://pubchem.ncbi.nlm.nih.gov/) and saved in SDF format. Files were converted from SDF to PDB format using UCSF Chimera software.

### Molecular docking

To conduct molecular docking, we selected the binding site of crystal ligands complexed with their reference protein and they are considered as a binding cavity.

Virtual screening was carried out using AutoDock Vina (AutoDock Vina 1.1.2 https://vina.scripps.edu) and the best ligand/ protein mode was identified based on the binding energy (kcal mol^-1^ units).In addition, are the same proteins are performed by Autdock 4 Suite tool.

### Docking sites by Autodock Vina with Pyrx Tool program

- *Structure of Sodium/glucose cotransporter 2* are: x = 38.7667663436 in Angstrom; y = 51.6876239274 in Angstrom; z = 45.0128531042 in Angstrom, with size X : 14.619903096 in Angstrom, size y = 17.3224718738 in Angstrom, size z = 17.6449783828 in Angstrom.
- *Structure of Sodium/glucose cotransporter 1* are: x = 99.9932915712 in Angstrom; y = 110.766137404 in Angstrom; z = 111.182435541in Angstrom, with size X : 25.0 in Angstrom, size y = 25.0 in Angstrom, size z = 25.0in Angstrom.

### Docking sites by Autodock 4 with MGL Tool program

- Center Grid Box of *NAD-dependent protein deacetylase SIRT1* are: x = 31.648 in Angstrom; y = -15.057 in Angstrom; z = 9.136 in Angstrom, with Spacing Grid Box: 0.375 Å, Population Size : 150, Number of energy evaluations: 2500000, num.grid points X (40), Y (40) Z(40).
- Center Grid Box of *NAD-dependent protein deacetylase SIRT2* are: x = 6.752 in Angstrom; y = -9.098 in Angstrom; z = -9.098 in Angstrom, with Spacing Grid Box: 0.375 Å, Population Size : 150, Number of energy evaluations: 2500000,num.grid points X (40), Y (40) Z(40).
- Center Grid Box of *NAD-dependent protein deacetylase SIRT3* are: x = 10.253in Angstrom; y = 3.097 in Angstrom; z = 5.237 in Angstrom, with Spacing Grid Box: 0.375 Å, Population Size : 150, Number of energy evaluations: 2500000,num.grid points X (40), Y (40) Z(40).
- Center Grid Box of *NAD-dependent protein deacetylase SIRT4* are: x = 18.565 in Angstrom; y = 25.607in Angstrom; z = -13.057 in Angstrom, with Spacing Grid Box: 0.375 Å, Population Size : 150, Number of energy evaluations: 2500000,num.grid points X (40), Y (40) Z(40).
- Center Grid Box of *NAD-dependent protein deacetylase SIRT5* are: x = 9.447 in Angstrom; y = -14.319 in Angstrom; z = 9.907in Angstrom, with Spacing Grid Box: 0.375 Å, Population Size : 150, Number of energy evaluations: 2500000,num.grid points X (40), Y (40) Z(40).
- Center Grid Box of *NAD-dependent protein deacetylase SIRT6* are: x = 22.911 in Angstrom; y = 2.572 in Angstrom; z = 7.454 in Angstrom, with Spacing Grid Box: 0.375 Å, Population Size : 150, Number of energy evaluations: 2500000,num.grid points X (40), Y (40) Z(40).
- Center Grid Boxof *NAD-dependent protein deacetylase SIRT7* are: x = 1.779 in Angstrom; y = 17.874 in Angstrom; z = 17.462 in Angstrom, with Spacing Grid Box: 0.375 Å, Population Size : 150, Number of energy evaluations: 2500000,num.grid points X (40), Y (40) Z(40).

## 3. Discussion and Results

The first class of proteins discussed in this Bioinformatic work was the Human Sodium/glucose cotransporter 2 (hSGLT2) and Human Sodium/glucose cotransporter 1 (SGLT1) cotransporter involved in the reabsorption of glucose in the kidney.

Particular attention, we focused on hSGLT2 that it mediates the reabsorption of the majority of filtered glucose in the kidney ^12,14^. Several studies in the literature reported that pharmacological inhibition of hSGLT2 by oral small-molecule inhibitors, such as empagliflozin, leads to enhanced glucose excretion and is widely used in the clinic to manage blood glucose levels for the treatment of type 2 diabetes (T2DM)^12,14^. SGLT2 inhibitors are a class of medications that modulate sodium-glucose transport proteins in the nephron (the functional units of the kidney), unlike SGLT1 inhibitors that perform a similar function in the intestinal mucosa. Two reviews have concluded that SGLT2 inhibitors benefit patients with atherosclerotic major adverse cardiac events (MACE)^12,14^. For these reasons, we performed several investigations by Molecular Docking of Polydatin with hSGLT2 and hSGLT1 to if this natural substance has good Binding Energy in the active pocket of these proteins. Several studies in the literature demonstrated that Polydatin, called Piceid, protects the heart from ischemic damage and it can be used for treating atherosclerotic diseases. ^4,5^

Indeed, in Fig. 1 is possible to observe the 3D structure of Sodium/glucose cotransporter 2 with a comparison pristine crystal ligand with best pose redocked same crystal ligand empagliflozin, called 73R with a binding energy value of -11.68 kcal/mol. In fact from docking analysis with Autodock Vina in the Ligand Binding Site pocket we obtained quite a perfect overlapping between two ligands (crystal and docked same ligand), with an RMSD (root-mean-square deviation) value of about 0.93 Ångström, showing that Molecular Docking with Autodock 4 with parameters of Grid box X, Y, and Z coordinates of this ligand 73R) could bring excellent Binding energies values of our target molecule (Polydatin) with Sodium/glucose cotransporter 2 (PDB Code: 7VSI).

**Fig 1.**
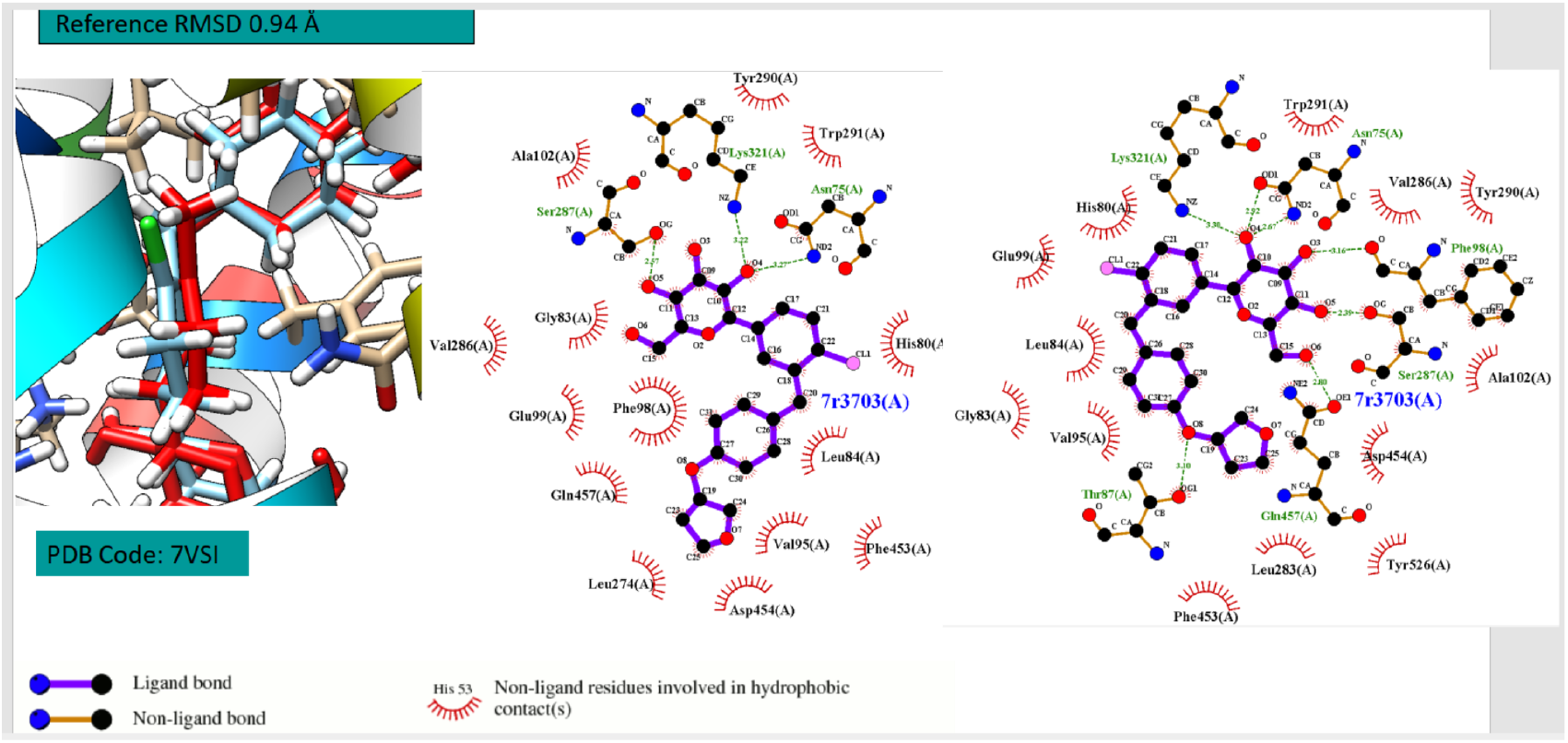
Comparison of overlapping of blu color pristine crystal ligand with red color 1 pose docked crystal ligand empagliflozin, called 73R : -11.68 kcal/mol (on the left panel); Comparison of 2D diagram residues interactions of pristine crystal ligand with red color 1 pose docked crystal ligand complexed with Structure of Sodium/glucose cotransporter 2 (on the right panel). The figure was reproduced by LigPlot program.

In addition, we showed the comparison of 2D diagram residues interactions bonds of pristine crystal ligand 73R with the best pose same ligand 73R complexed with hSGLT2, identifying with LigPlot software which are the hydrogen bonds and hydrophobic bonds that are established between the ligand 73R and human SGLT2 protein. In Fig. 2, we conducted the same analysis and characterization of Polydatin, after performing its docking analysis by Autodock 4 by MGL tool. In Fig. 3 is possible to observe the 3D structure of Sodium/glucose cotransporter 1 with a comparison pristine crystal ligand with best pose redocked same crystal ligand 1YI with a binding energy value of -12.12 kcal/mol. In fact from docking analysis with Autodock Vina in the Ligand Binding Site pocket we obtained quite perfect overlapping between two ligands (crystal and docked same ligand), RMSD (root-mean-square deviation) value of about 1.43 Ångström, showing that Molecular Docking with Autodock 4 with parameters of Grid box X, Y, and Z coordinates of this ligand 1YI) could bring excellent Binding energies values of our target molecule (Polydatin) with Sodium/glucose cotransporter 2 (PDB Code: 7WMV). In Fig.4 we reported the same docking analysis and characterizations focused on Polydatin. In short, the main docking results of hSGLT2 and hSGLT1 were reported in Table 1, where it is possible to infer that apparently, the Polydatin is capable of doing well with both hSGLT1 and hSGLT2.

**Fig 2.**
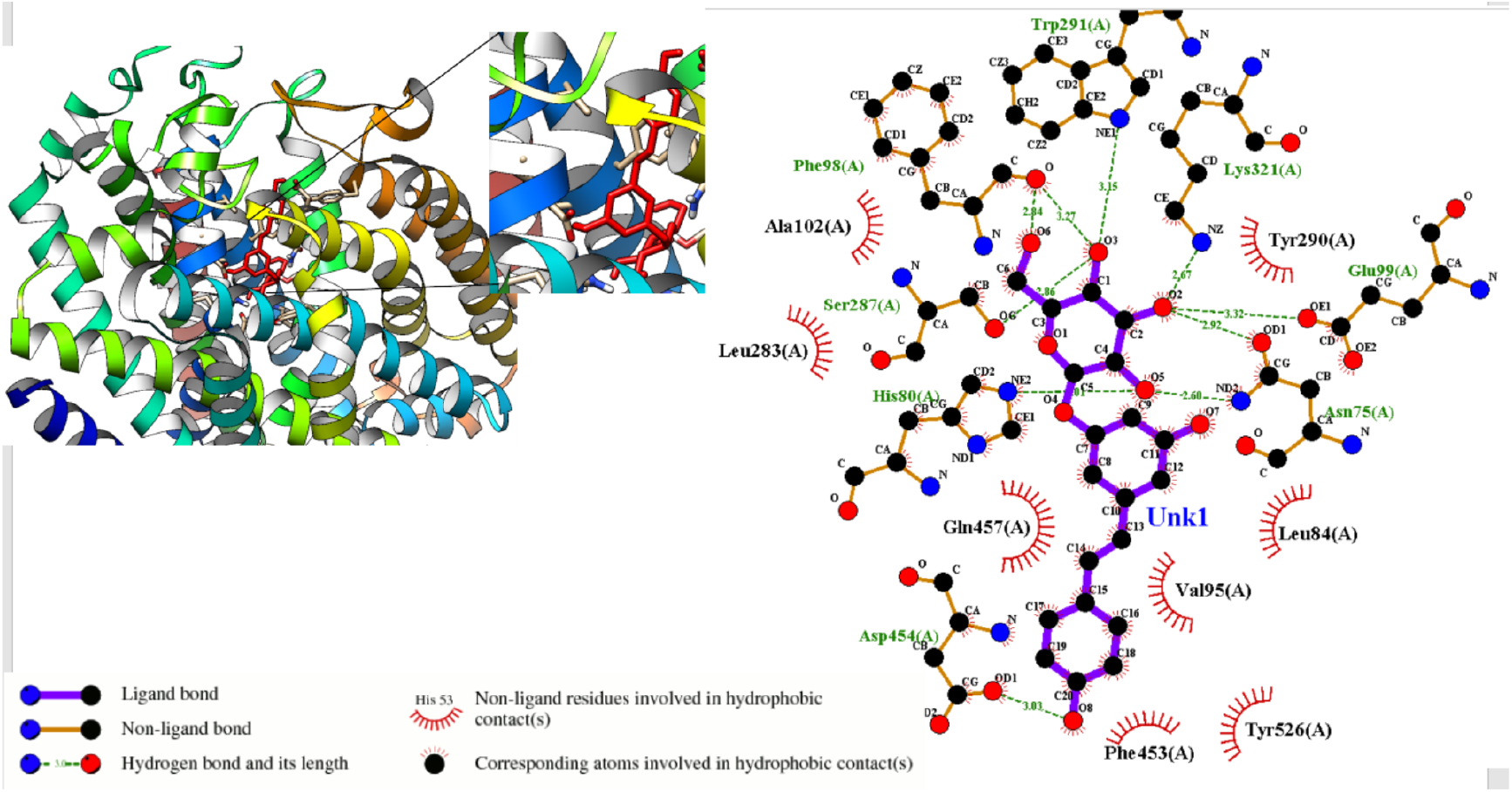
3D structure of Structure of Sodium/glucose cotransporter 2 with complexed docked Polydatin : -9.82 kcal/mol (on the left side); 2D diagram residues interactions of docked Polydatin. The figure was reproduced by LigPlot program.

**Fig 3.**
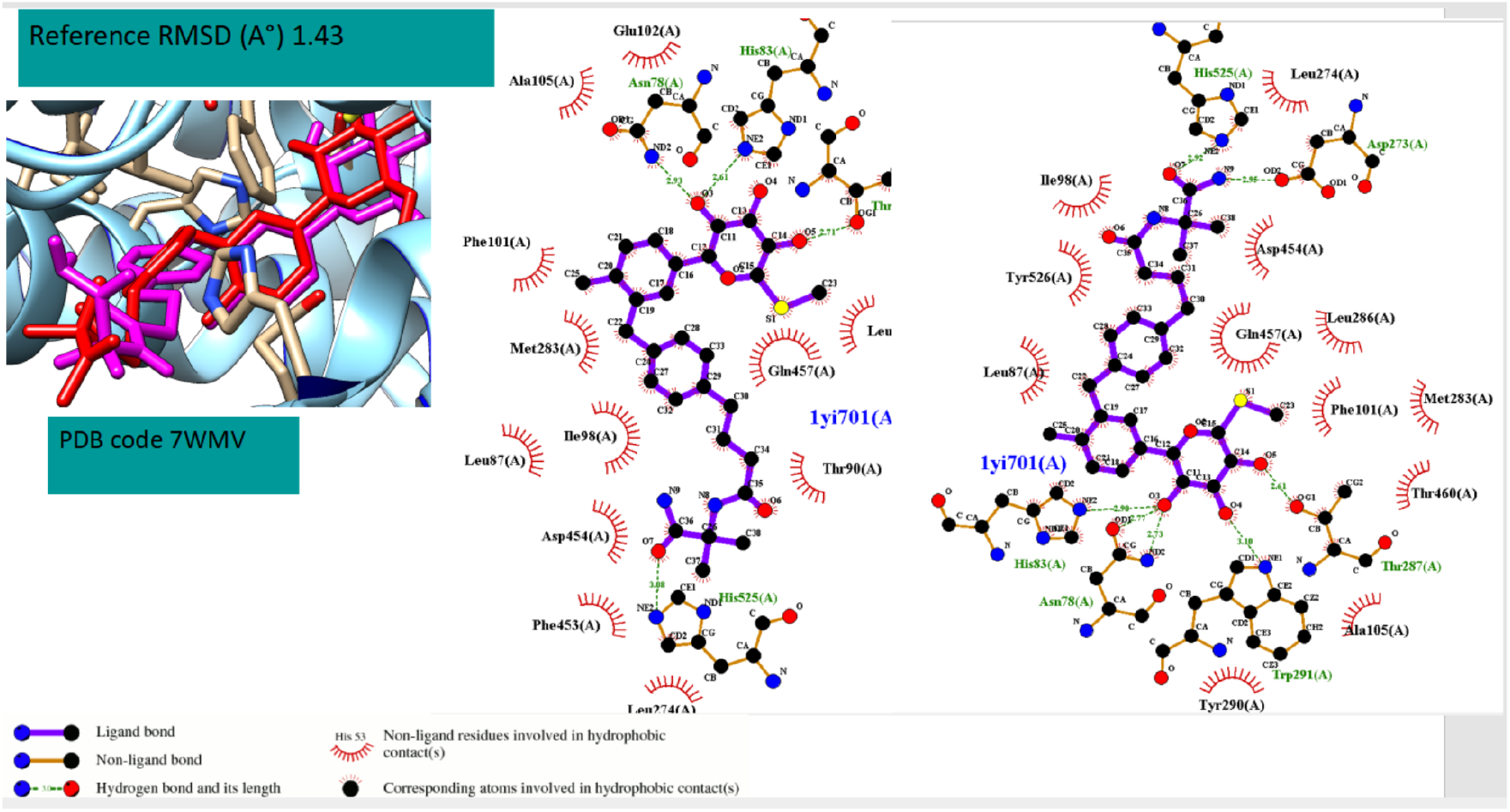
Comparison of overlapping of blu color pristine crystal ligand with red color 1 pose docked crystal ligand 1YI : -12.12 kcal/mol (on the left panel); Comparison of 2D diagram residues interactions of pristine crystal ligand 1YI with red color 1 pose docked crystal ligand complexed with Structure of Sodium/glucose cotransporter 1 (on the right panel). The figure was reproduced by LigPlot program.

**Fig 4.**
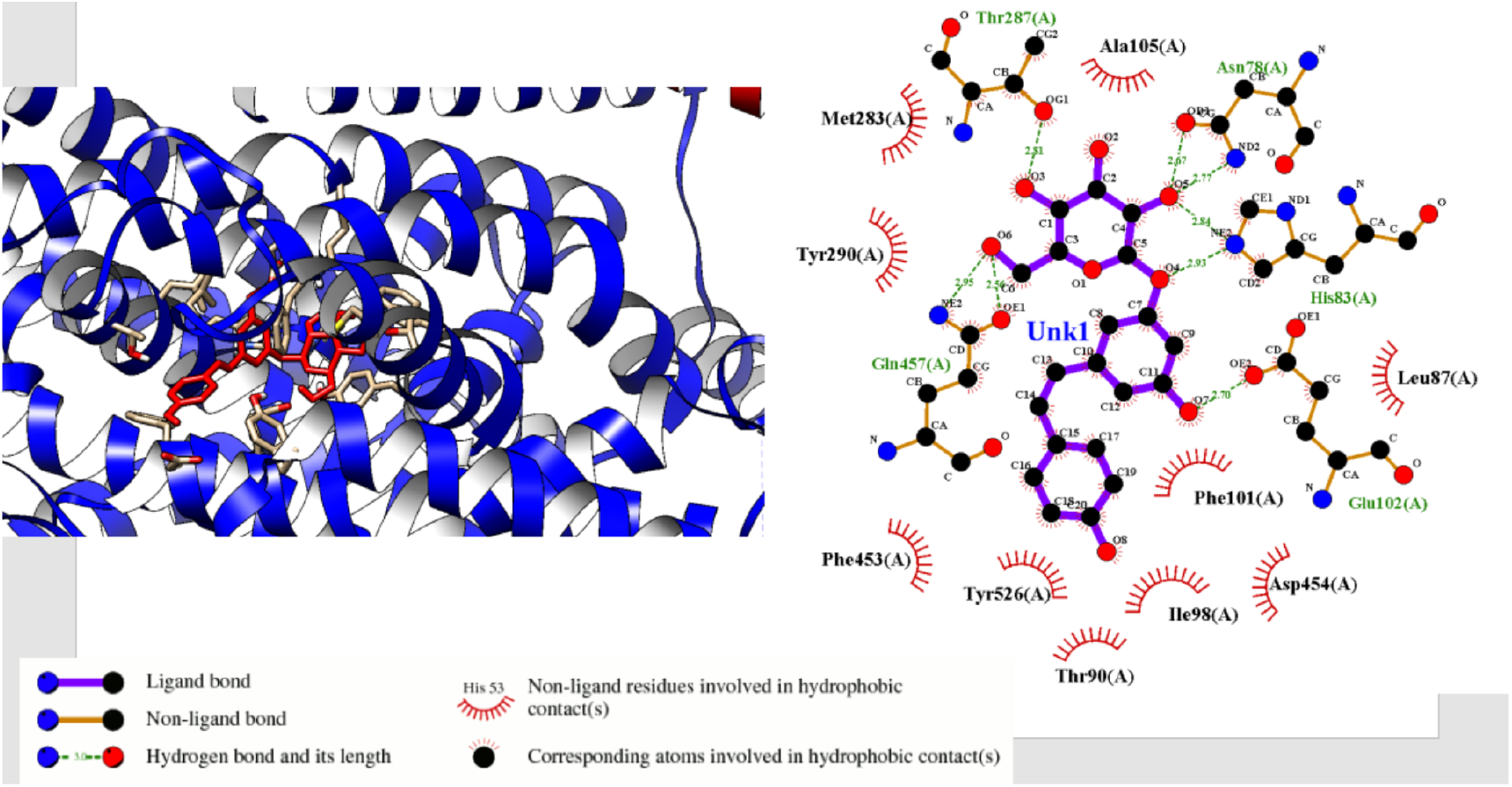
3D structure of Structure of Sodium/glucose cotransporter 1 with complexed docked Polyda-tin : -9.29kcal/mol (on the left side); 2D diagram residues interactions of docked Polydatin. The figure was reproduced by LigPlot program.

**Table 1.** Comparison Docking results by Autodock Vina and Autodock 4 of Structure of Sodium/glucose cotransporter 1 and Structure of Sodium/glucose cotransporter 2 with complexed crystal ligand 73R and 1YI respectively and with docked Polydatin.

| Compounds | PDB Code | Binding Energy by Vina<br>Score (kcal/mol) | Binding Energy by<br>Autodock 4 (kcal/mol) |
| --- | --- | --- | --- |
| Crystal Ligand° 1YI | 7WMV | -11,10 | -12,12 |
| Polydatin | 7WMV | -10,00 | -9,25 |
| Crystal Ligand 73R | 7VSI | -11,70 | -11,68 |
| Polydatin | 7VSI | -10,00 | -9,82 |

The second class of human proteins discussed in this communication was SIRTs Family named Sirtuins, involved in several important biological processes. They are proteins that help us regulate homeostasis, the condition of cells and their regeneration, support metabolism, keep the brain healthy, and make you age healthily. Sirtuins, a family of nicotinamide adenine dinucleotide (NAD +)-dependent enzymes affect mitochondria and aging. They have enzymatic activity by functioning as a histone deacetylase or mono-ribosyl transferase. Functioning as metabolic sensors, sirtuins, and longevity proteins, play a fundamental role in maintaining the integrity of the genome, as well as participating in the maintenance of the normal state of chromatin condensation. ^15-21^.

Above all, we want to underline one of their important aspects, that of protecting the heart. In fact, according to some studies they are involved in the processes of atherosclerosis 17-21. For all these functions, we decided to investigate Polydatin for the first time, by Molecular Docking both Autodock Vina and Autodock 4 Algorithms with all seven sirtuin proteins, to understand where it binds, and with whom among them it has an excellent ability to bind on active site of these proteins.

From Fig. 5-11, we report 2D diagram residues interactions reproduced by the Ligplot program of Structure of Human NAD-dependent protein deacetylase sirtuin-1, sirtuin2,sirtuin-3,sirtuin-5,sirtuin-6,sirtuin-7, with their Crystal ligand, their redocked same ligand and docked Polydatin in Ligand BindingSite pocket, except for the Sirtuin 4 which originated from Xenopus tropicalis. The 2D analysis obtained by Ligplot allows you to automatically calculate what types of bonds can be formed with the investigated protein. In fact, by carefully observing the figures above reported, it is possible to identify both the hydrogen bonds (green in color) and the hydrophobic bonds involved in this protein investigated (indicated with semicircles).

**Fig 5.**
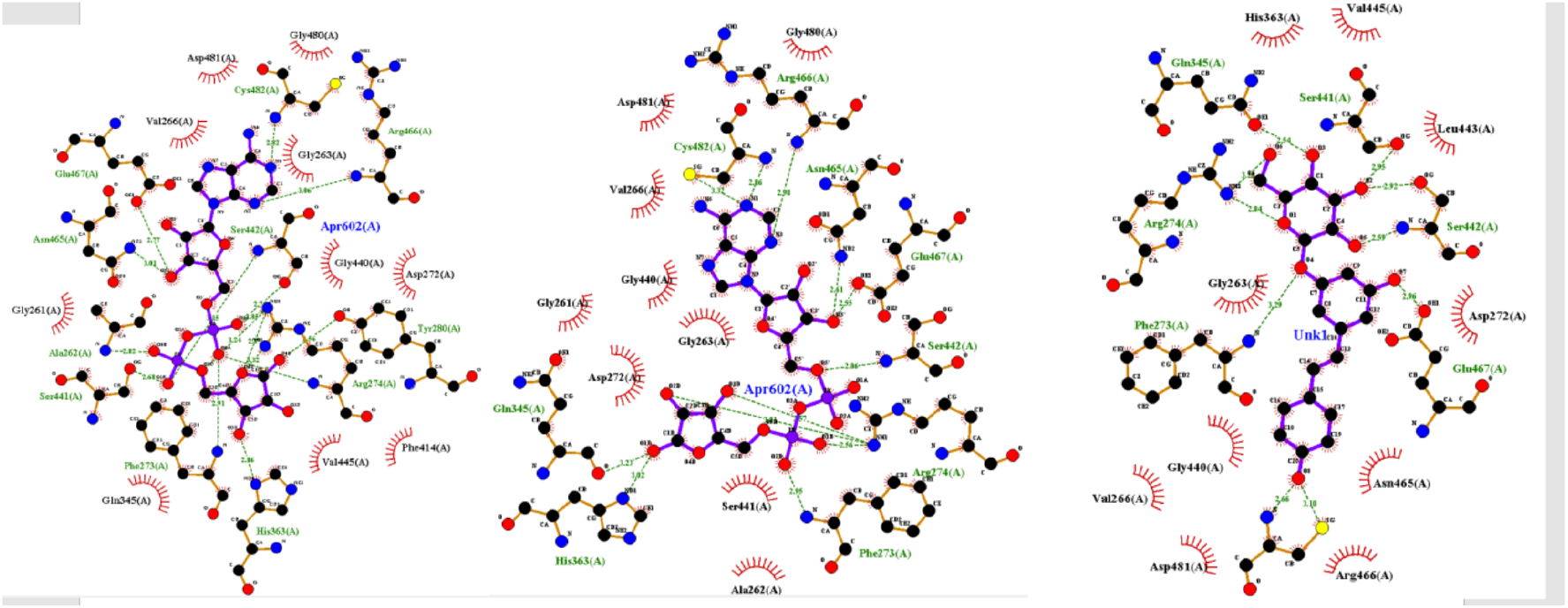
2D diagram residues interactions bonds of Structure of Human NAD-dependent protein deacetylase sirtuin-1 with Crystal ligand, docked same ligand APR -9.7 kcal/mol and docked Polydatin -10.47 kcal/mol, in Ligand Binding Site pocket (PDB 4KXQ). The figure was reproduced by Lig-Plot program. **\***Crystal ligand Adenosine-5-Diphosphoribose called APR

**Fig 6.**
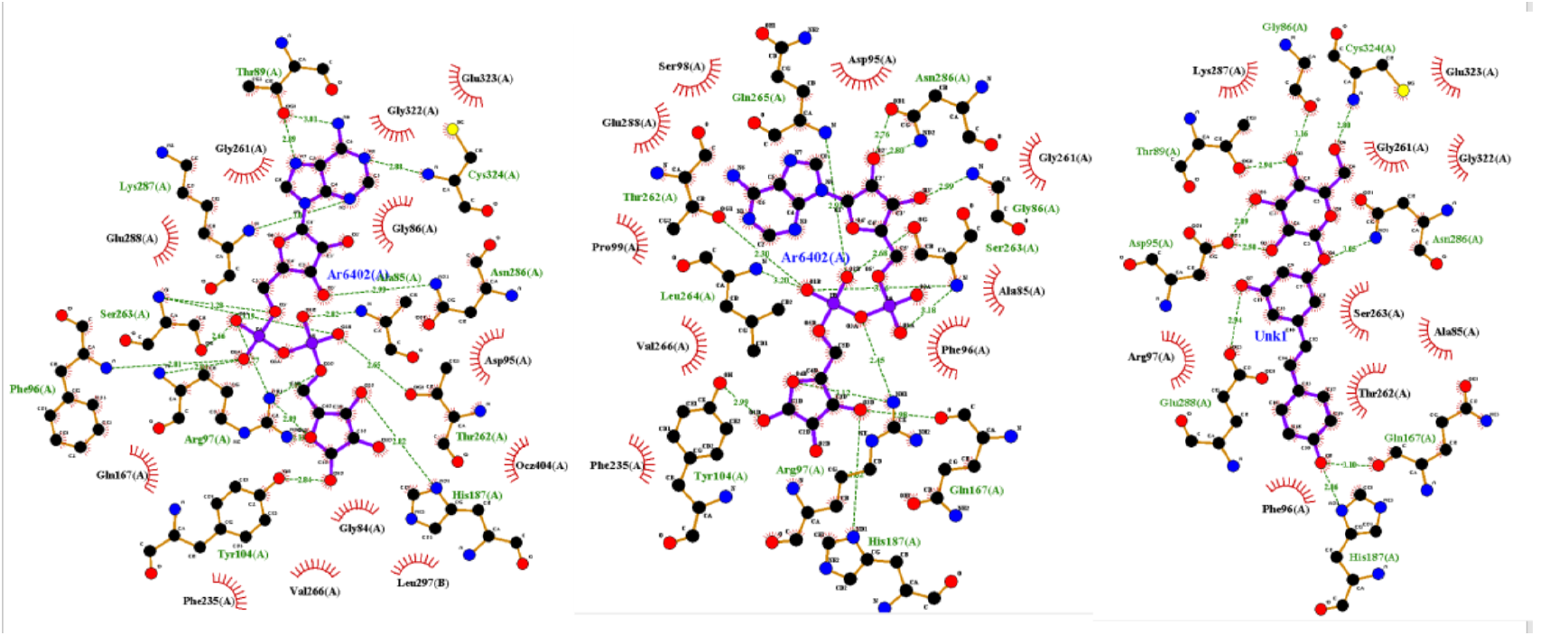
2D diagram residues interactions bonds of Structure of Human NAD-dependent protein deacetylase sirtuin-2 with Crystal ligand* Adenosine-5-Diphosphoribose called APR, docked same ligand APR -7.43 kcal/mol and docked Polydatin -10.49 kcal/mol, in Ligand Binding Site pocket (PDB 5D7P). The figure was reproduced by LigPlot program. **\***Crystal Ligand AR6 : [(2R,3S,4R,5R)-5-(6-AMINOPURIN-9-YL)-3,4-DIHYDROXY-OXOLAN-2-YL]METHYL [HYDROXY-[[(2R,3S,4R,5S)-3,4,5-TRIHYDROXYOXOLAN-2-YL]METHOXY]PHOSPHORYL] HYDROGEN PHOSPHATE

**Fig 7.**
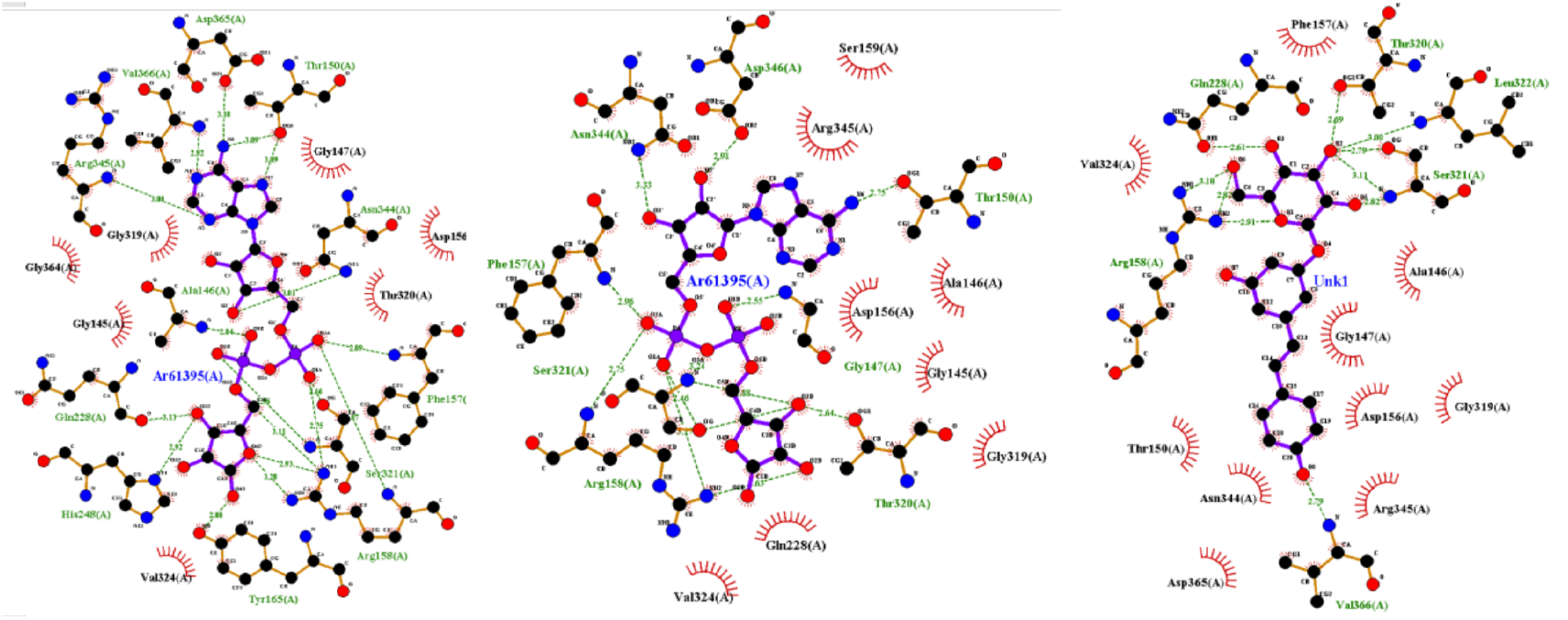
2D diagram residues interactions bonds of Structure of Human NAD-dependent protein deacetylase sirtuin-3 with Crystal ligand, docked same ligand -9.32 kcal/mol and docked Polydatin -10.43 kcal/mol, in Ligand Binding Site pocket (PDB 4BN4). The figure was reproduced by Lig-Plot program. **\***Crystal Ligand AR6 : [(2R,3S,4R,5R)-5-(6-AMINOPURIN-9-YL)-3,4-DIHYDROXY-OXOLAN-2-YL]METHYL [HYDROXY-[[(2R,3S,4R,5S)-3,4,5-TRIHYDROXYOXOLAN-2-YL]METHOXY]PHOSPHORYL] HYDROGEN PHOSPHATE

**Fig 8.**
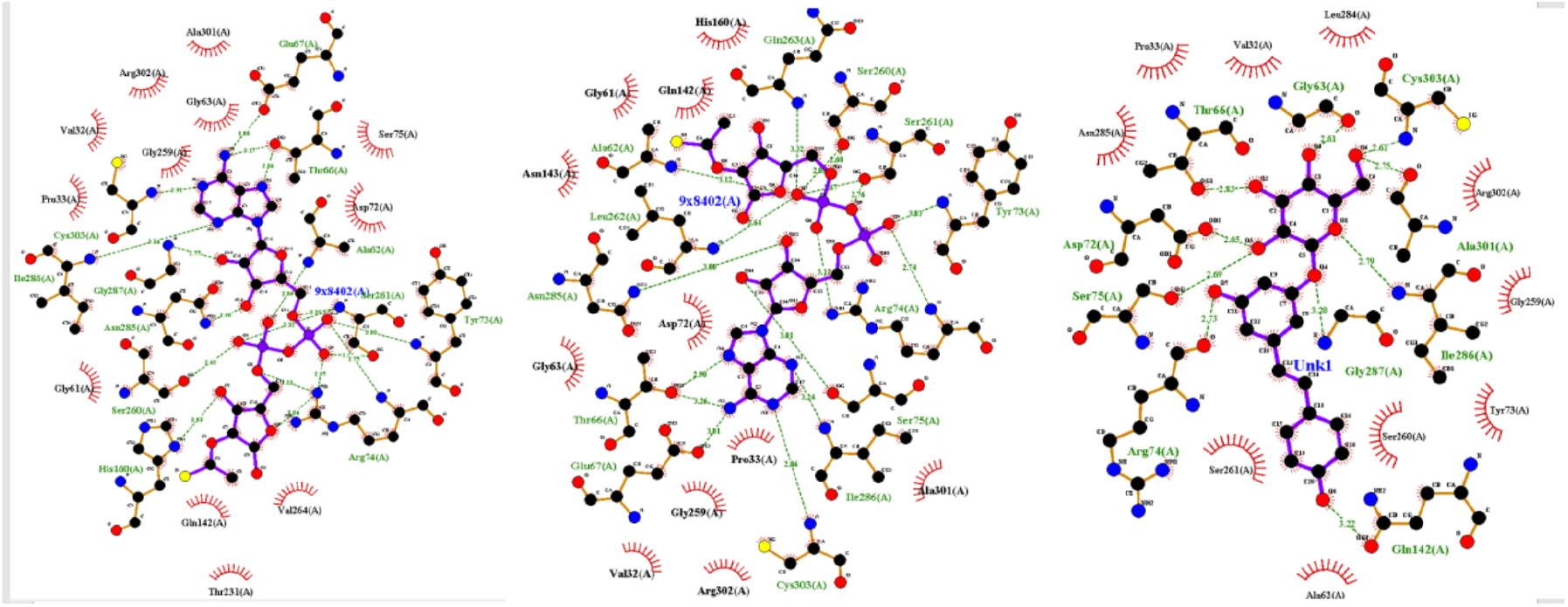
2D diagram residues interactions bonds of Structure of NAD-dependent protein deacetylase sirtuin-4 with Crystal ligand Adenosine-5-Diphosphoribose called APR, docked same ligand APR -12.81 kcal /moland docked Polydatin -11.71 kcal/mol, in Ligand Binding Site pocket (PDB 5OJN). The figure was reproduced by LigPlot program. **\*** Crystal ligand 9X8 thioacetyl-ADP-ribose; Sirtuin 4 Chain A originated from Xenopus tropicalis

**Fig 9.**
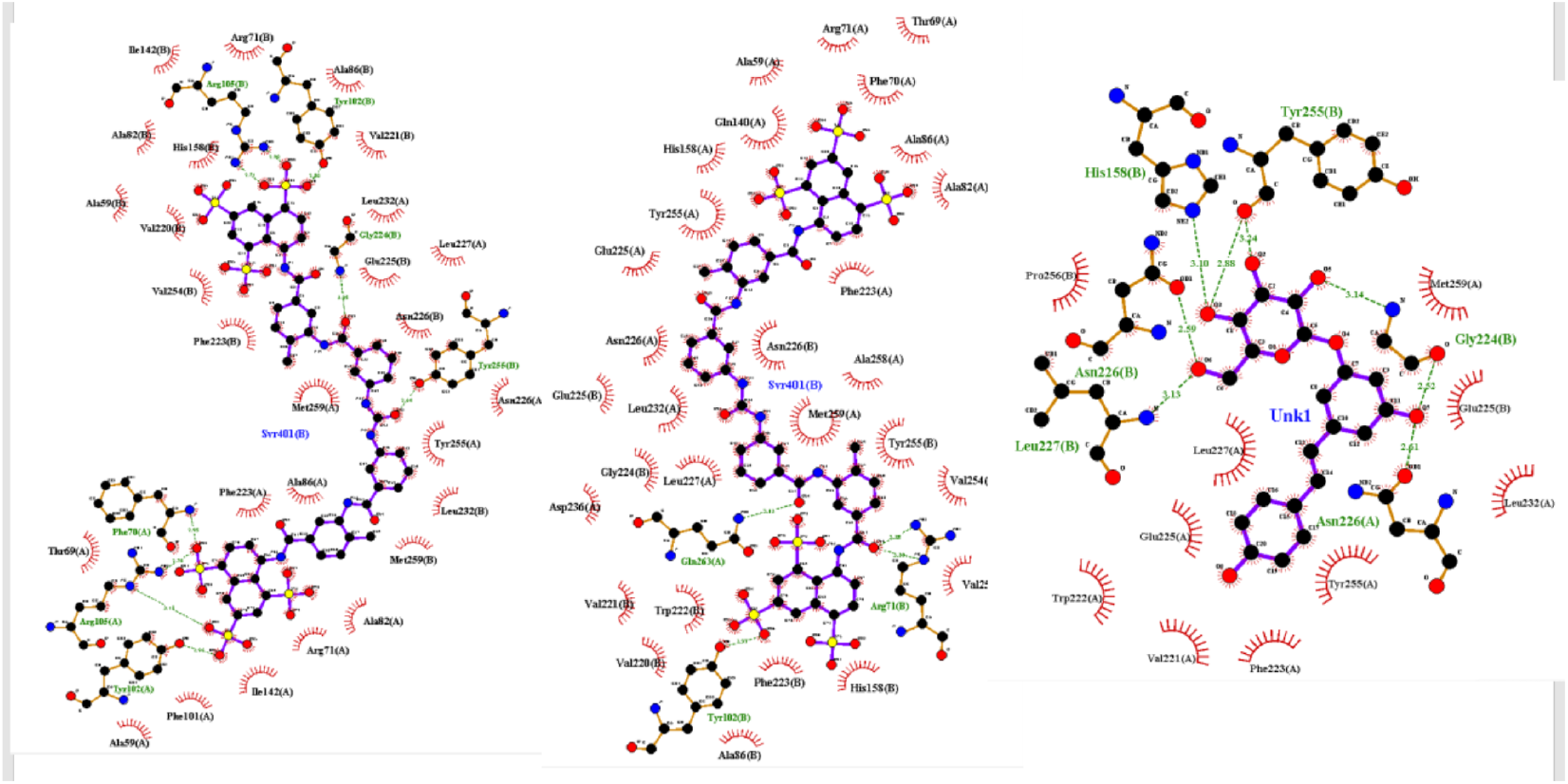
2D diagram residues interactions bonds of Structure of Human NAD-dependent protein deacetylase sirtuin-5 with Crystal ligand, docked same ligand *12.82 kcal/mol APR and docked Polydatin -10.05 kcal/mol, in Ligand Binding Site pocket (PDB 2NYR). The figure was reproduced by Lig-Plot program. **\***Crystal Ligand SVR 8,8’-[CARBONYLBIS[IMINO-3,1-PHENYLENECARBONYLIMINO(4-METHYL-3,1 PHENYLENE)CARBONYLIMINO]]BIS-1,3,5-NAPHTHALENETRISULFON IC ACID

**Fig 10.**
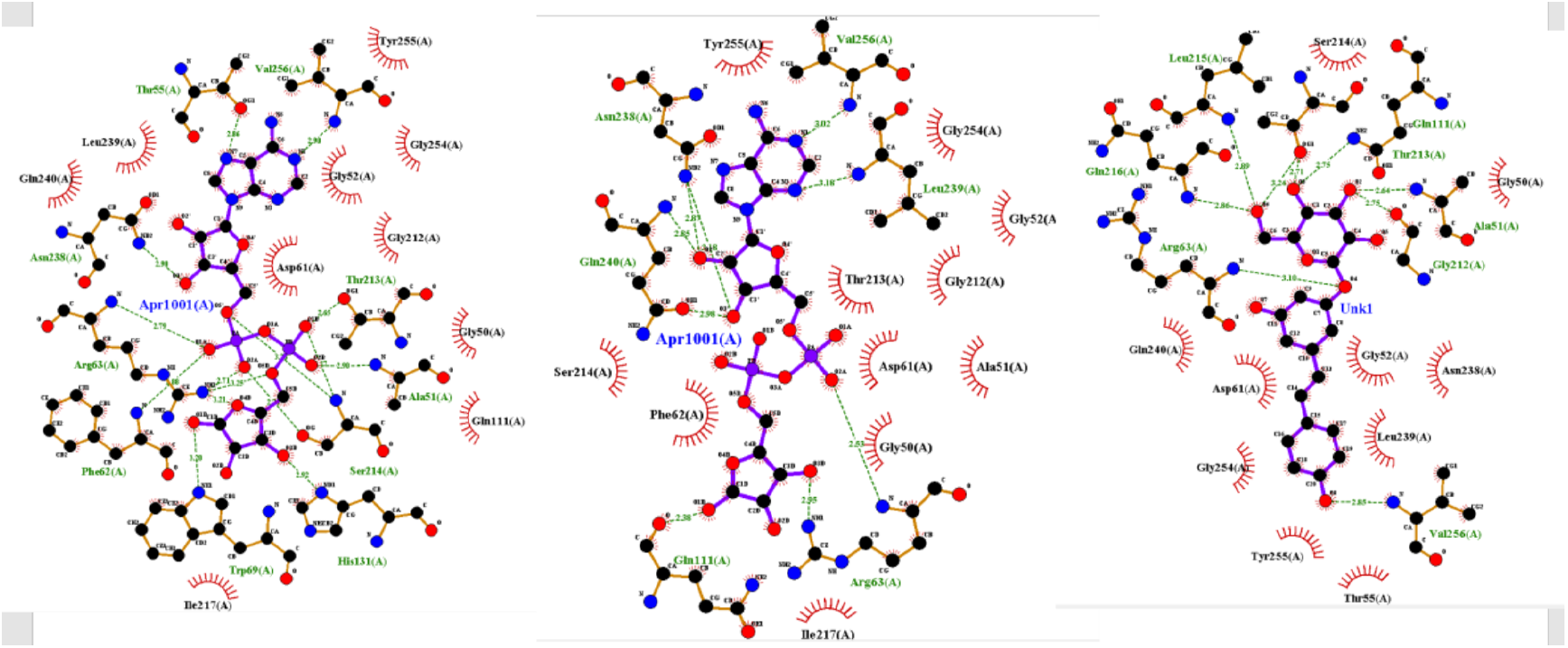
2D diagram residues interactions bonds of Structure of Human NAD-dependent protein deacetylase sirtuin-6 with Crystal ligand, docked same ligand -8.52 kcal /mol and docked Polydatin -10.72 kcal/mol, in Ligand Binding Site pocket (PDB 3K35). The figure was reproduced by Lig-Plot program. **\***Crystal ligand Adenosine-5-Diphosphoribose called APR

**Fig 11.**
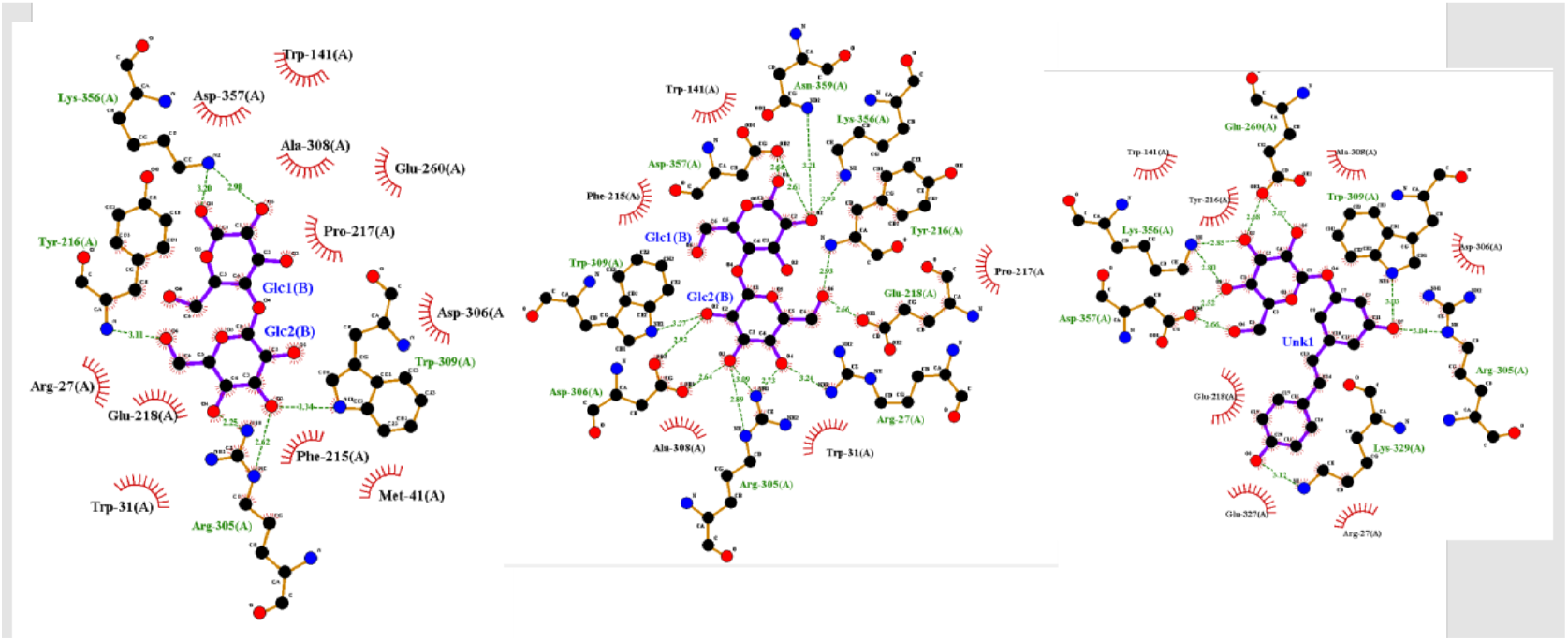
2D diagram residues interactions bonds of Structure of Human NAD-dependent protein deacetylase sirtuin-7 with Crystal ligand docked same ligand -6.78 kcal/mol and docked Polydatin -9.17 kcal/mol, in Ligand Binding Site pocket (PDB 5IQZ). The figure was reproduced by LigPlot program. **\***Crystal Ligand alpha-maltose

Fig. 12 showed the theoretical role of Polydatin with its excellent ability to bind to the whole Sirtuin family at the N terminal (SIRT1-SIRT-7), showing Binding Energies scores of about -10/11 kcal/mol, significant values if compared to the respective ligand crystals, which have only a value of about -9/-10 kcal/mol.

**Fig 12.**
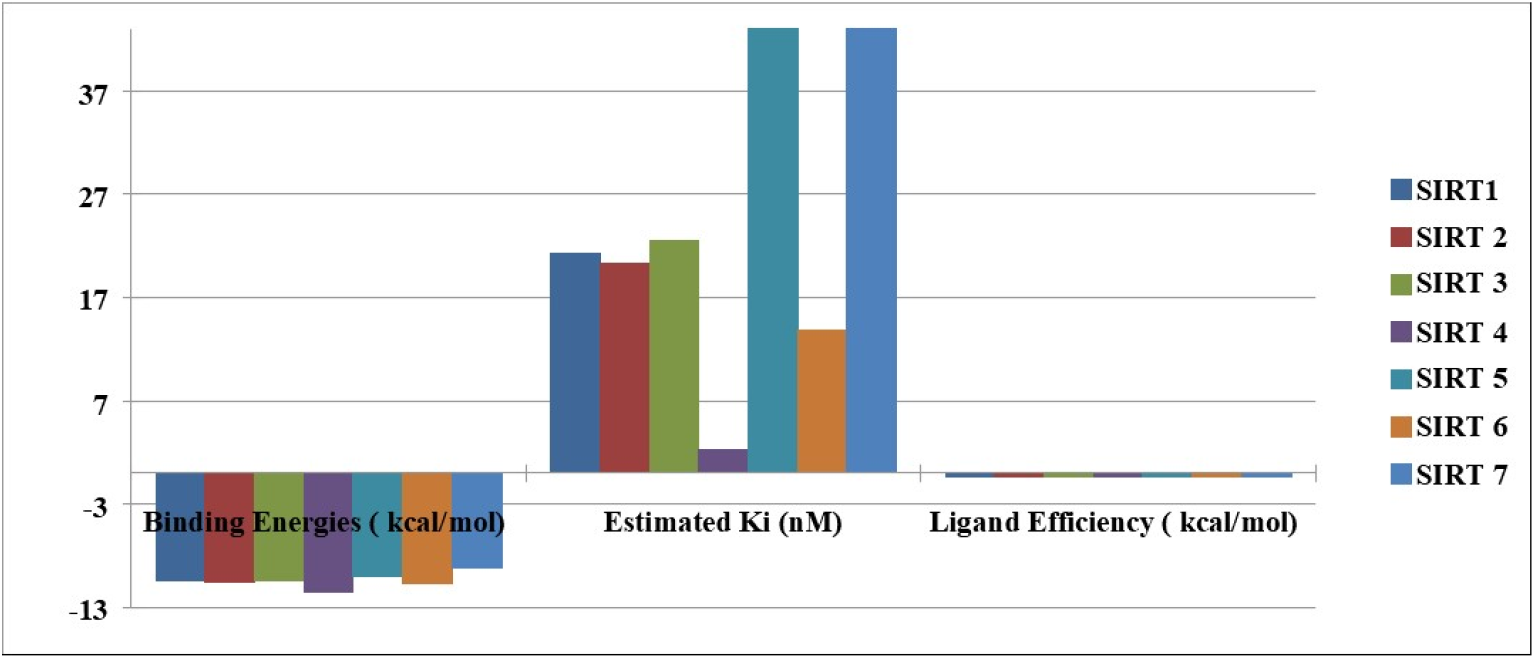
Comparison Docking results by Autodock 4 (Binding energies scores, Estimaiton constant inhbtions Ki and ligand efficiencies of Structure of SIRTs proteins with docked Polydatin.

In addition, if we carefully observe the predicted inhibition constants ki scores (in Nanomolar units), we find out significant differences rather than those binding energies values. In fact, with the same Binding energies scores, Polydatin has been shown to possess a low Ki value with Sirtuin 6, with a value of about 13 nMolar, and with Sirtuin 4, with a value of about 2 nMolar. With human sirtuins 1,2,3 Polydatin had a value of about 20 nMolar. Concerning sirtuin 5 has a higher value of about 40 nMolar and sirtuin 7 about 190 nMolar.

Unfortunately, the value of sirtuin 4, even though is very good does not refer to human sirtuin but to Xenopus Tropicalis, because a version of the aforementioned human protein in a crystallized form satisfactory to be studied by Molecular Docking is not available from the PDB Protein database.

## 4. Conclusions

One important aspect we underlined is that our in silico analyses are accurate even though are necessary experiments in vitro, in vivo, and clinical studies to confirm a real biological function of polydatin both human SGLT2 and human SGLT1 and beyond with human Sirtuins proteins. However, Molecular Docking can be a useful tool to understand its chemical-physical behavior with the proteins investigated with the Piceid in this work. From all the computational results, Polydatin has shown excellent results potentially proving to be a natural medicine capable of possessing excellent biological properties with high binding energies and low constant inhibitions Ki (in nanoMolar units) with both proteins class (SGLT1/SGLT2)and (SIRTs family) discussed in this paper.

## References

1. Beerling, D. J., Bailey, J. P., & Conolly, A. P. (1994). Fallopia japonica (Houtt.) ronse decraene. Journal of Ecology, 82(4), 959–979.

2. Shang, Y., Zhang, P., Wei, W., Li, J., & Ye, B. C. (2023). Metabolic engineering for the high-yield production of polydatin in Yarrowia lipolytica. Bioresource Technology, 129129.

3. Du, Q. H., Peng, C., & Zhang, H. (2013). Polydatin: a review of pharmacology and pharmacokinetics. Pharmaceutical biology, 51(11), 1347–1354.

4. Luo, S. F., Yu, C. L., & Zhang, P. W. (1990). Influences of 3, 4, 5-trihydroxystibene-3-beta-mono-D-glucoside on beat rate and injury of cultured newborn rat myocardial cells. Zhongguo yao li xue bao= Acta Pharmacologica Sinica, 11(2), 147–150.

5. San Tang, K., & Tan, J. S. (2019). The protective mechanisms of polydatin in cerebral ischemia. European journal of pharmacology, 842, 133–138.

6. Cheng, Y., Zhang, H. T., Sun, L., Guo, S., Ouyang, S., Zhang, Y., & Xu, J. (2006). Involvement of cell adhesion molecules in polydatin protection of brain tissues from ischemia–reperfusion injury. Brain research, 1110(1), 193–200.

7. Lanzilli, G., Cottarelli, A., Nicotera, G., Guida, S., Ravagnan, G., & Fuggetta, M. P. (2012). Anti-inflammatory effect of resveratrol and polydatin by in vitro IL-17 modulation. Inflammation, 35, 240–248.

8. Wu, M., Li, X., Wang, S., Yang, S., Zhao, R., Xing, Y., & Liu, L. (2020). Polydatin for treating atherosclerotic diseases: a functional and mechanistic overview. Biomedicine & Pharmacotherapy, 128, 110308..

9. Jakhar, R., Dangi, M., Khichi, A., & Chhillar, A. K. (2020). Relevance of molecular docking studies in drug designing. Current Bioinformatics, 15(4), 270–278.

10. Trott, O., & Olson, A. J. (2010). AutoDock Vina: improving the speed and accuracy of docking with a new scoring function, efficient optimization, and multithreading. Journal of computational chemistry, 31(2), 455–461.

11. Rizvi, S. M. D., Shakil, S., & Haneef, M. (2013). A simple click by click protocol to perform docking: AutoDock 4.2 made easy for non-bioinformaticians. EXCLI journal, 12, 831.

12. Zelniker, T. A., Wiviott, S. D., Raz, I., Im, K., Goodrich, E. L., Bonaca, M. P., … & Sabatine, M. S. (2019). SGLT2 inhibitors for primary and secondary prevention of cardiovascular and renal outcomes in type 2 diabetes: a systematic review and meta-analysis of cardiovascular outcome trials. The Lancet, 393(10166), 31–39.

13. Xu, D., Chandler, O., Wee, C., Ho, C., Affandi, J. S., Yang, D., … & Xiao, H. (2021). Sodium-Glucose Cotransporter-2 Inhibitor (SGLT2i) as a Primary Preventative Agent in the Healthy Individual: A Need of a Future Randomised Clinical Trial?. Frontiers in Medicine, 8, 712671.

14. Niu, Y., Liu, R., Guan, C., Zhang, Y., Chen, Z., Hoerer, S., … & Chen, L. (2022). Structural basis of inhibition of the human SGLT2–MAP17 glucose transporter. Nature, 601(7892), 280–284.

15. Houtkooper, R. H., Pirinen, E., & Auwerx, J. (2012). Sirtuins as regulators of metabolism and healthspan. Nature reviews Molecular cell biology, 13(4), 225–238.

16. Covarrubias, A. J., Perrone, R., Grozio, A., & Verdin, E. (2021). NAD+ metabolism and its roles in cellular processes during ageing. Nature Reviews Molecular Cell Biology, 22(2), 119–141.

17. Tanno, M., Kuno, A., Horio, Y., & Miura, T. (2012). Emerging beneficial roles of sirtuins in heart failure. Basic research in cardiology, 107, 1–14.

18. Matsushima, S., & Sadoshima, J. (2015). The role of sirtuins in cardiac disease. American Journal of Physiology-Heart and Circulatory Physiology, 309(9), H1375–H1389.

19. Bugger, H., Witt, C. N., & Bode, C. (2016). Mitochondrial sirtuins in the heart. Heart Failure Reviews, 21(5), 519–528.

20. Dolinsky, V. W., Cole, L. K., Sparagna, G. C., & Hatch, G. M. (2016). Cardiac mitochondrial energy metabolism in heart failure: Role of cardiolipin and sirtuins. Biochimica et Biophysica Acta (BBA)-Molecular and Cell Biology of Lipids, 1861(10), 1544–1554.

21. Yang, X., Chang, H. C., Tatekoshi, Y., Balibegloo, M., Wu, R., Chen, C., … & Ardehali, H. (2023). SIRT2 inhibition protects against cardiac hypertrophy and heart failure. bioRxiv, 2023-01.

